# Unveiling theranostic potential: Insights into cell-free microRNA-protein interactions

**DOI:** 10.1101/2024.12.23.630210

**Authors:** Vishal Kumar Sahu, Subhayan Sur, Sanjana Agarwal, Harish Madhyastha, Amit Ranjan, Soumya Basu

**Affiliations:** Cancer and Translational Research Centre, Dr. D. Y. Patil Biotechnology and Bioinformatics Institute, Dr. D. Y. Patil Vidyapeeth, Tathawade, Pune - 411033, India; Department of Cardiovascular Physiology, Faculty of Medicine, University of Miyazaki, Miyazaki 8891692, Japan

**Keywords:** Cell-free miRNA, miRNA-protein interaction, miRNA-protein docking, Oncomirs, miRNA-protein database

## Abstract

MicroRNAs (miRNAs) belong to a short endogenous class of non-coding RNAs which have been well studied for their crucial role in regulating cellular homeostasis. Their role in modulation of diverse biological pathways by intracellular or extracellular communication and interaction with DNA, RNA or protein, projects them or their targets as promising biomarkers and therapeutic agents. However, studying specific interactions in the extracellular or cell-free environment for drug discovery or biomarker establishment is resource-intensive. In this study, we derive a computational approach based on available experimental data to decipher patterns in miRNA-protein interactions in the cell-free milieu. We characterized the miRNA-protein interactome (miRPin, https://www.mirna.in/miRPin) and identified consensus sequences governing these interactions. The study establishes the role of multiple miRNAs and protein interactions present in cell-free environments leading to pathological conditions, viz., role of proteins like METTL3 and AGO2 etc. and miRNAs like hsa-miR-484 and hsa-miR-30 families, hsa-mir-126-5p etc. in association with multiple miRNAs in different cancer types, cardiovascular diseases, and neurological disorders. The findings outlined in the study may facilitate new avenues of therapeutic discovery leading to the understanding of cellular mechanisms underlying therapy relapse and drug resistance. By addressing these interactions in an extracellular environment may further gain insight into regulating disease initiation and progression, overcoming challenges related to drug efficacy and drug delivery in the presence of bilayer membranes with drug efflux pumps.

## 1. Introduction

Discovery of microRNAs (miRNAs) has changed the purview of understanding of disease mechanisms and since then their interaction has been widely explored with other RNAs, genes, and proteins within the cells and in the cell-free environment such as blood plasma and different body fluids [1–4]. In the extracellular environment, miRNAs are usually found to be associated with extracellular vesicles (EVs) like microvesicles and exosomes resulting in intercellular communications [5,6]. However, detection of miRNA molecules in unprotected and ribonuclease (RNase) rich environments, such as saliva, urine, serum, or blood plasma is gaining attention for their theranostic potential alongside the speculations on the stability of RNA molecules in many types of body fluids due to the presence of RNases [7–10]. This requires developing deep insight into the stability and role of RNA molecules specially miRNAs in body fluids and cellular environment.

It has been more than a decade since miRNA and protein complexes were identified in the extracellular environment and realized for their diagnostic and therapeutic potential. Firstly, Wang et. al. reported binding of Nucleophosmin (NPM1) and other nucleolins with miRNAs like miR-122, -671-3p, and -943 in cell culture setup [3]. Argonaute complexes (AGO1 or AGO2) were soon reported to be binding with miRNAs (miR-142–3p, miR-122, miR-24, and miR-150) in blood plasma [11,12]. Additionally high-density lipoproteins have been identified to be binding with different miRNAs such as miR-125a, miR-223, miR-24, miR-27, miR-146, miR-222 along with their uptake via class B type I scavenger receptors [6,13]. Demonstration of uptake of AGO2-miRNA complexes into the cells was a significant advancement towards the understanding of the complete lifecycle of miRNAs through their secretion and delivery to target cells [14]. However, the mechanism and specific role of primary or secondary structure of interacting miRNA-protein partners and their varying existence in circulation is poorly understood in relation with different stages of diseases.

Apart from normal expression of cell-free miRNAs (cf-miRNAs) or proteins, these biomolecules have been widely explored for their expression in the context of different diseases such as neurological disorders, cardiovascular diseases, disorders of the immune system, inflammatory diseases, and cancers etc. Cancers based on its origin and subtype show differential expression of miRNA and proteins extending the pattern in body fluids as well regulating different hallmarks of the cancer [15–17]. Differential expression of miRNAs in cells and serum have been studied in different types of neurological diseases including Parkinson’s, age-related neurodegenerative diseases, Alzhiemer’s, and Schizophrenia linking to different stages of disease progression [2,18–21]. Multiple serum miRNAs have been reported to be associated with different diabetes related diseases or glucose metabolism related disorders [22–24]. Although information on coexpression of miRNA-protein, miRNA-mRNA, miRNA and other non-coding RNAs is available, however, information on structure based interactions of miRNA-protein specific to cell-free environment is sparsely available [25–27].

There exist advanced and promising approaches to predict the miRNA-protein interactions using computational and laboratory methods [28,29]. However, their limited exploitation and advancement leaves a void of understanding of structure based miRNA binding with protein and their synergistic role. Hence, constructing a data-intensive landscape of cell-free miRNAs-protein interactome and their binding patterns using computational study can be of paramount significance for deciphering the precise mechanisms underlying disease initiation and progression. Investigating the interplay between multimodal miRNA-protein interactomes may offer valuable insights into factors contributing to therapy relapse, drug inefficacy, and drug resistance, thereby paving the way for alternative, advanced, and effective therapeutic strategies.

This study explores sequence specific interactions of extracellular proteins with cf-miRNAs and implications of these interactions. The prediction of miRNA-protein interacting patterns will hold promise for precision therapeutic and diagnostic strategies enabling accurate disease staging and treatment especially in cancer. This might advance our understanding of cancer progression and treatment as well.

## 2. Methods

### 2.2. Curation of miRNA-protein interactome in extracellular environment

Publicly available resources containing experimental evidence of miRNA-protein interactions were utilized for the preparation of miRNA-protein interactome data set. The list of proteins expressed in extracellular vesicles and plasma was obtained using the Plasma Proteome Database (PPD) and Vesiclepedia. PPD contains information of proteins detected in serum or plasma using mass spectrometry [30]. Vesiclepedia is a manually curated source for the biomolecules present in subtypes of extracellular vesicles (EVs) including exosomes, microvesicles, migrasomes, apoptotic bodies, and large oncosomes [31]. The miRBase database, a primary repository for miRNA information, was employed to get a list of all miRNAs [32]. The gene names of miRNAs were mapped using HUGO Gene Nomenclature Committee (HGNC) approved identifiers for miRNAs and their genes [33]. Circulating microRNA Expression Profiling (CMEP) database and miRNATissueAtlas2 database were explored to annotate miRBase miRNAs as extracellular miRNAs [34,35].

These enlisted extracellular proteins were further marked for RNA binding domains (RBDs) and miRNA-protein associations using Cross Linked immunoprecipitation sequence database (CLIPdb) and RBPbase [36,37]. The CLIPdb database contains experimentally deduced miRNA-protein associations involving cross-linked immunoprecipitation of RNAs with antibodies for protein [36]. CLIPdb had miRNA identifiers to validate miRNA co-precipitated proteins providing experimental evidence for miRNA-protein binding. The RBPbase database by ‘The European Molecular Biology Laboratory (EMBL)’ contains annotations of RNA binding domains in proteins for humans (*Homo sapiens*) as well as other species [37]. Proteins annotated with RBDs in humans were selected for this study.

MiRTarBase is a database of experimentally verified miRNA and target interactions (MTIs) using relevant methods such as western blotting, microarrays, or qPCR [38]. The targets of miRNAs were obtained from miRTarBase. Further the interactions were filtered and mapped with the aforementioned miRNA and protein gene lists.The dataset served as starting point to study extracellular or cell-free interactome of miRNA and protein.

### 2.3. Sequence and structure information of miRNAs and proteins

miRBase was used for obtaining sequence information of miRNAs [32]. To obtain structural information, we utilized a structure database miRVim, which contains predicted and experimentally available three-dimensional structures of miRNAs [39]. Gene identifiers or official gene names for the proteins were submitted to UniProt Knowledgebase (UniProtKB) to obtain sequence and structure information for proteins [40]. The protein structures were obtained from the RCSB-PDB database and AlphaFoldDB [41,42]. AlphaFoldDB was utilized to retrieve predicted protein structures in case of incomplete structure or unavailability of the structure in the RCSB repository.

### 2.4. Gene set enrichment analysis of cell-free the proteins and miRNAs

The genes for protein and miRNA were mapped with their official gene symbols standardized by HUGO Gene Nomenclature Committee (HGNC) [33]. These HGNC identifiers were submitted to DAVID server for gene set enrichment analysis (GSEA) to identify major role, localisation, or significant molecular function asserted by the miRNAs and proteins [43]. DAVID is a web server for annotation, visualization and integrated discovery of provided genes to study their involvement and functional role in various biological pathways by associating the genes with gene ontology (GO) terms. The enrichment results of GO terms with *p*-value less than 0.05 and false discovery rate (FDR) less than 0.1 (<10%) were considered significant in the GSEA results.

### 2.5. Discovery of patterns among interacting entities

The cluster of proteins interacting with miRNAs and the cluster of miRNAs interacting with proteins were prepared and the sequence information of the grouped entities was stored in FASTA file format for motif discovery. This was to identify possible common patterns in miRNAs sequences interacting with a protein and *vice-versa*.

The motif discovery in the grouped protein and miRNA sequences were conducted using MEME-suite, a set of tools to thoroughly investigate sequences under study [44]. XSTREME, a comprehensive motif analysis tool, conducted in-depth analyses, including motif discovery, on sequences where motif sites could be positioned anywhere [45]. XSTREME accommodated input sequences of variable lengths leveraging various tools like Sensitive, Thorough, Rapid, Enriched Motif Elicitation (STREME), Multiple EM for Motif Elicitation (MEME), Simple Enrichment Analysis (SEA), Motif Comparison Tool (Tomtom), and Find Individual Motif Occurrences (FIMO), for seamless pattern discovery and sequence enrichment analysis [46–50].

MEME played a crucial role in identifying novel and ungapped motifs characterized by recurring fixed-length patterns within the sequences. Additionally, MEME efficiently partitioned variable-length patterns into distinct motifs for further analysis. STREME focused on discovering enriched motifs by comparing their prevalence in the sequences to control sequences. Tomtom facilitated motif comparison against known motifs in databases, namely PROSITE (a database of protein families and domains) and RNA binding motif database [51,52]. It ranked motifs and produced alignments for significant matches. Tomtom was provided with MEME recognisable motif dataset from PROSITE (MEME Reference: PROTEIN/prosite2021_04.meme) for the protein sequences and RBP (MEME Reference: RNA/Ray2013_rbp_Homo_sapiens.meme) for microRNA sequences [51,52]. SEA was utilized to determine the most enriched motifs to get the insights into motif enrichment. FIMO played a vital role in scanning sequences to find individual matches with the specified motifs, aiding in identifying occurrences of motifs within the sequences. Together, these tools offered extensive insights into motif discovery and characterization within the analyzed sequences.

The cutoff of Expectation-value calculated by XSTREME program was set to less than or equal to 0.05 (*E*-value ≤ 0.05) for the statistical significance of the predicted and enriched motifs.

### 2.6. Molecular docking study to rank protein-RNA motif interactions

Interacting miRNA and protein pairs were combined for the discovered motifs for docking analysis. The aim was to find the interactions between the discovered motifs and to explore the possible precise sequence of interaction between the miRNAs and proteins. The discovered motifs were extracted from the three-dimensional structure of the proteins and miRNAs using ChimeraX and Visual Molecular Dynamics (VMD) packages using their available scripting interface [53,54]. These extracted structures were docked and scored using HDock Lite to find protein-RNA interactions by performing protein-nucleic acid docking study [55].

The ranks for interactions were calculated by assigning rank based weights to the score, number of contacts, and hydrogen bonds of the predicted docking conformations. These weights were further refined with the composite index construction method [56]. In the context of protein-RNA interaction analysis, composite index construction involves integrating multiple variables like scores, number of contacts, hydrogen bonds, solvent accessible surface area (SASA) of the receptor fragment, SASA of ligand fragment, and distance between the geometric centers of the fragments obtained from docking analysis into a single metric for ranking the conformations. The advantage of using a composite index is that it allows for a more comprehensive evaluation of interactions compared to relying on individual criteria alone. It takes into account the relative importance of each factor. This approach helps prioritize and identify the most significant protein-RNA interacting conformers.

Bulk data processing and queries were performed using python programming and scripting language.

## 3. Results and Discussion

### 3.1. miRNA-protein interactome in extracellular vesicles and blood plasma

A miRNA-protein interactome (miRPin) was curated by leveraging information available in public databases containing experimental evidence of localisation of miRNAs and proteins and their interactions. Relative information available in the databases were correlated to compile a list of miRNAs and proteins having interaction in cell-free environment (Figure 1).

**Figure 1:**
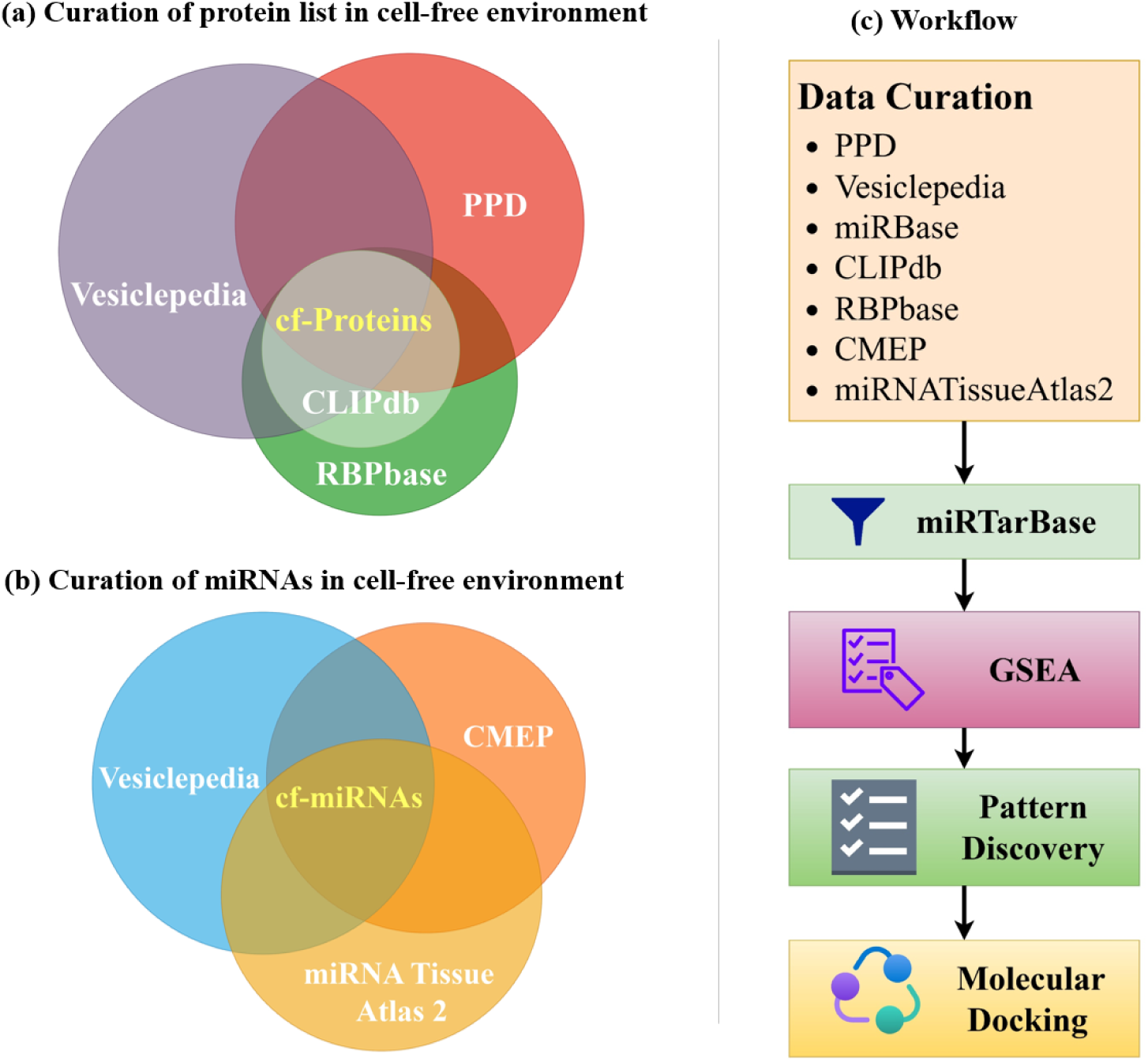
(a) Annotation of cell-free proteins was done by finding overlapping proteins in Vesiclepedia, PPD, CLIPdb and RBPbase databases. Similarly, the list of miRNAs was curated from Vesiclepedia, CMEP, and miRNA Tissue Atlas 2 databases. (c) The list of miRNA and proteins were filtered in miRTarBase for their interactions.The information was further used for gene set enrichment analysis (GSEA), pattern discovery, and molecular docking. (PPD:; CMEP:

The Plasma Proteome Database (PPD) cataloged a total 10,422 proteins and Vesiclepedia contained information of 18,391 proteins. Further RNA binding domains and experimental evidence of binding with RNA in the protein list was identified using the CLIPdb and RBPbase database that contained 221 and 4,257 protein gene identifiers respectively. Correlation of CLIPdb and RBPbase with Vesiclepedia and PPD yielded 198 extracellular proteins identifying 158 proteins in extracellular and non-vesicular space while 192 in EVs (Supplementary File 1). The correlation approach helped in narrowing down proteins that bind with miRNAs fulfilling the criteria of having an RNA binding motif within their structures and have been reported to bind with RNA molecules. Among the extracellular proteins, 38 proteins were found to be exclusively present in the extracellular vesicle i.e., listed in Vesiclepedia while absent in PPD. Similarly, 4 proteins (YBX3, RTCB, LSM11, and AGO2) were exclusively present in the blood plasma.

EV miRNAs were validated with the miRNAs cataloged in Vesiclepedia. The processes helped culminate in the association of 2,163 miRNAs with plasma and 1,916 miRNAs with EVs (Supplementary File 2). The miRTarBase database yielded 21,060 unique miRNA-protein interactions out of a total of 957,040 experimentally validated interactions providing a complete overview of possible extracellular interaction dataset (Supplementary File 3). The compiled and annotated lists of extracellular or cell-free miRNAs and proteins (available at https://www.mirna.in/miRPin) are not exhaustive and serve as a foundational resource for further exploration of gene ontology, pattern discovery, and molecular docking in our study.

### 3.2. Proteins and miRNAs exhibited functional diversity in the terms of gene ontology

Gene set enrichment analysis (GSEA) performed using DAVID yielded a diverse landscape of gene ontology (GO) terms for the available 2,027 miRNA and 198 protein gene identifiers. Proteins exhibited enrichment across a broad spectrum of GO terms spanning Cellular Component (CC), Molecular Function (MF), and Biological Process (BP) (Supplementary File 4). In contrast, microRNAs displayed pronounced enrichment primarily in BP terms, indicating their involvement in various biological processes (Supplementary File 5). However, microRNAs showed comparatively lower enrichment in CC and MF categories. This differential enrichment suggests distinct functional roles and regulatory mechanisms governed by proteins and microRNAs in cellular processes and biological pathways.

#### 3.2.1 Cell-free miRNAs (cf-miRNAs) participate in diverse biological processes

The list of enriched GO terms for miRNAs provided possible involvement of the selected cf-miRNAs in different biological pathways however were limited in case of molecular function and localisation. Biological processes related to gene silencing, translation inhibition, and degradation of mRNA by miRNA were highly enriched (>29% miRNAs) (Figure 2a). Along with their broad regulatory roles, a wide-range of different biological processes like angiogenesis, apoptosis, cellular response to virus, ion transport, regulation of cardiac muscles, phenotypic switching, and regulation of cell adhesion etc. had lower frequency. The GO terms related to localisation were limited to RISC Complex, Extracellular Space, Extracellular Vesicle, and Extracellular Exosomes only (Figure 2b). Additionally, the selected cf-miRNAs were marked for only two molecular functions: Posttranscriptional gene silencing by binding with messenger RNA (mRNA) and Binding with mRNA at 3’ untranslated regions (Figure 2c).

**Figure 2:**
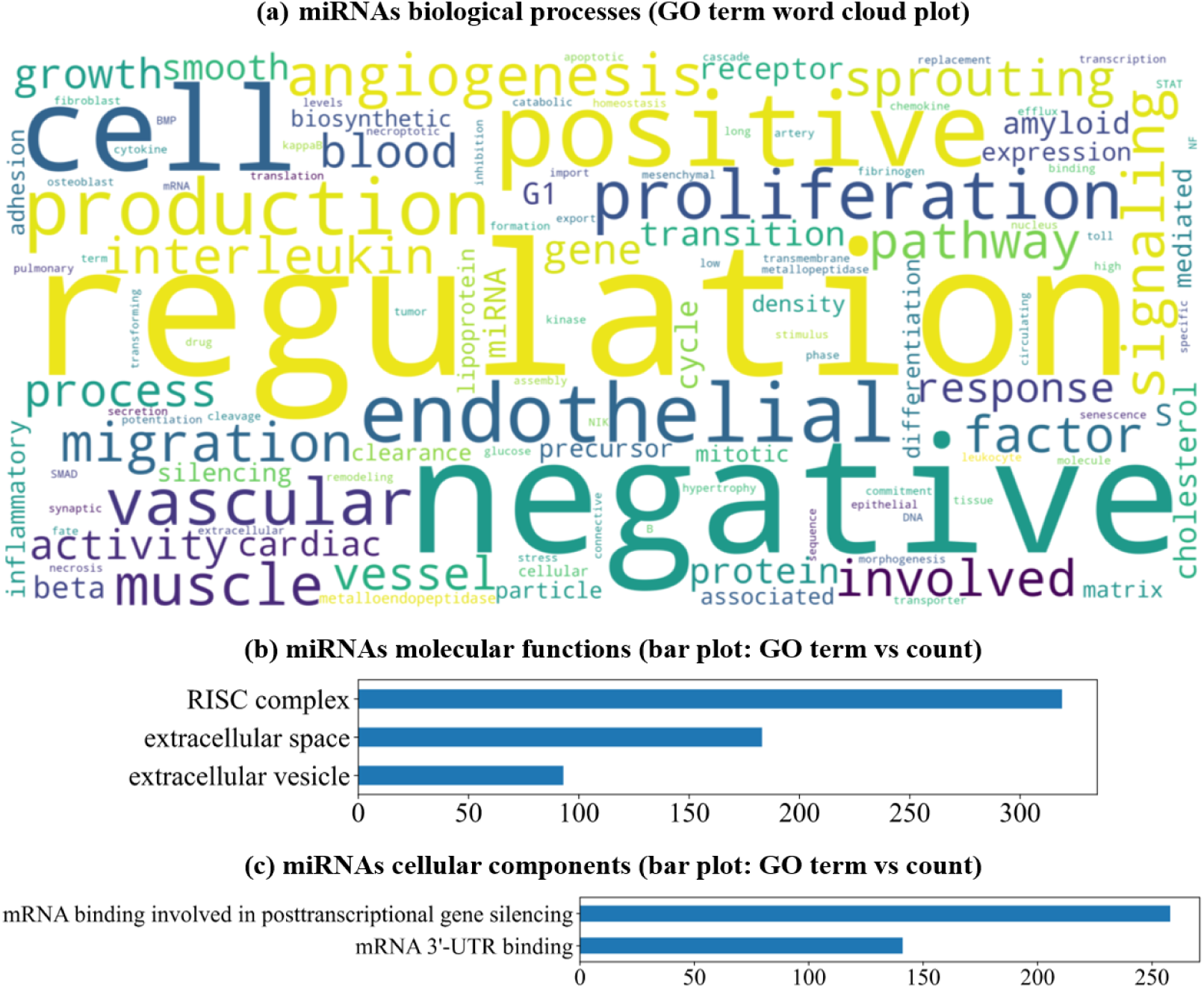
Gene Set Enrichment Analysis (GSEA) of miRNAs. (a) Word cloud plot of 53 unique terms; (b) cf-miRNAs MF GO terms enriched for only two molecular functions posttranscriptional gene silencing by binding with messenger RNA (mRNA) and binding with mRNA at 3’ untranslated regions; (c) Cellular components enrichment bar graph.

Around 70 BP terms out of 136 for miRNA GSEA were related to negative regulation of various processes while 30 BP terms were related to positive regulation (Figure 2a). The enrichment analysis confirmed the localisation of the selected miRNAs to be maximum in extracellular space yet their participation in normal processes and various pathological conditions that can be associated with cardiovascular diseases, neurological disorders, and cancer hallmarks. This signifies that the miRNA in the extracellular environment should to be explored for their ability for modulating the cellular response of the distant cells as a result of coordinated communication at different stages of pathological conditions (Figure 2).

#### 3.2.2 Cell-free proteins demonstrated distinctive functional heterogeneity

A total of 198 gene names were submitted to DAVID that yielded a pool of 86 BP, 47 CC, and 46 MF terms. The enrichment results of the biological processes were diverse and related to gene regulation assistance including RNA processing and transport (Figure 3a). The proteins were assorted to different compartments of the cell like nucleus, cytoplasm, synapse, Cajal body, P-body, membrane etc (Figure 3b). Additionally, they performed diversified molecular functions such as binding (binding with mRNA, RNA, or protein) and enzymatic activities (Figure 3c).

**Figure 3:**
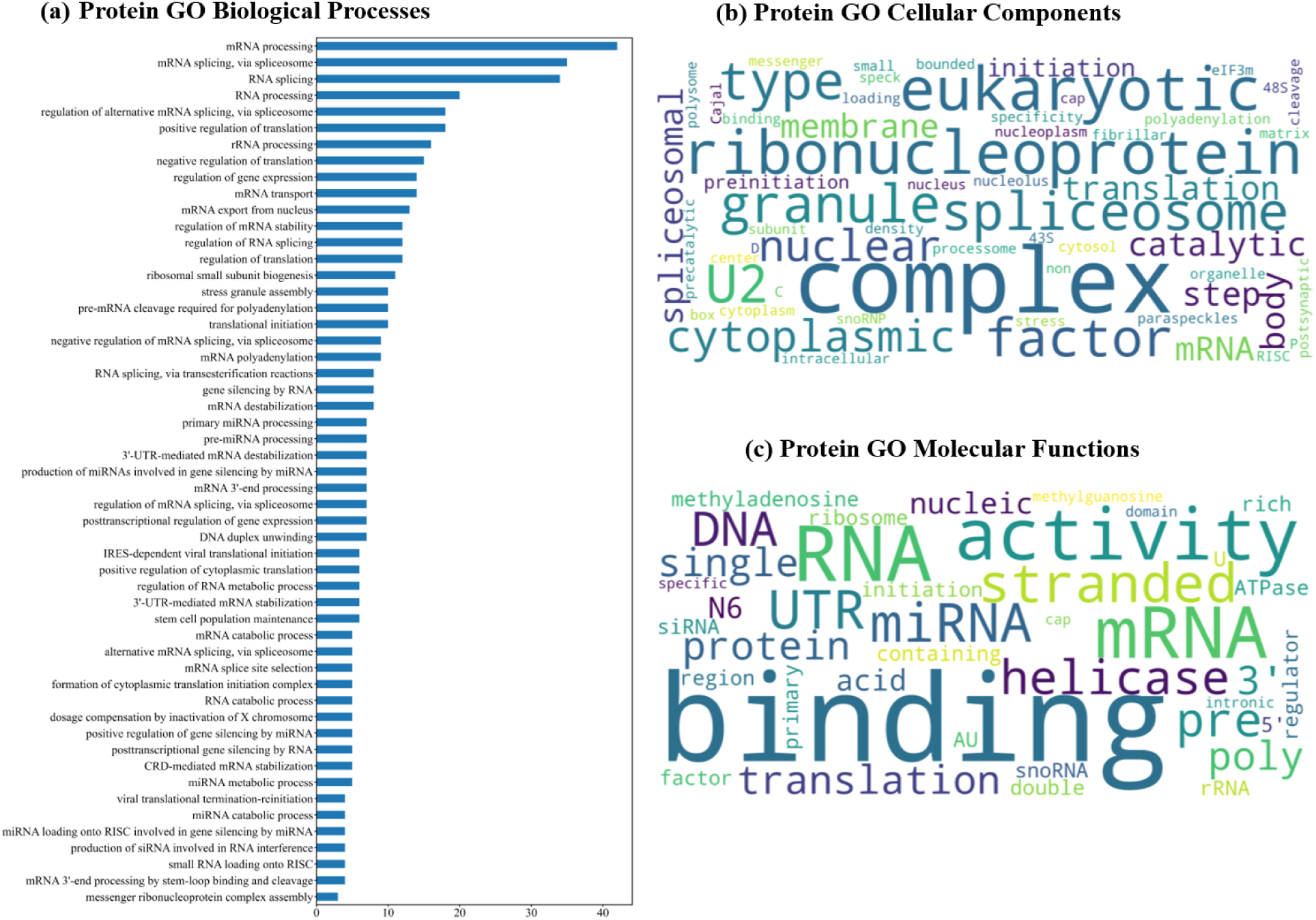
Protein genes enriched in diverse Gene Ontologies, (a) Proteins participated in distinctive regulatory processes in the cells while occupying different parts of the cellular location (b). Additionally, the molecular functions of the proteins are diverse which includes various enzymatic activities performed by the selected proteins (c).

Since proteins and miRNAs selected for cell-free environments participate in varied processes and activities, the proteins could be dispatching multifaceted roles including gene regulation, signal transduction, carrier for miRNAs in extracellular space, and protection from the RNase rich environment of the serum, blood plasma, saliva, and other body fluids [6,13]. This additionally strengthens our hypothesis and recommendation to explore the miRNA-protein complexes in an extracellular environment with relation to their life cycle in distant cellular communication.

### 3.3. MEME-suite assisted discovery of patterns in the sequences of cell-free miRNAs and proteins

Total 1,704 clusters for miRNAs (1,514) and proteins (192) were processed for pattern discovery. MEME suite analysis provided insight into the pool of motifs in miRNAs and proteins. XSTREME discovered a total of 43,614 motifs including miRNAs and proteins (Table 1 and Supplementary File 6). XSTREME discovered 2,874 consensus sequences for miRNAs while 940 sequences for proteins.

**Table 1:**
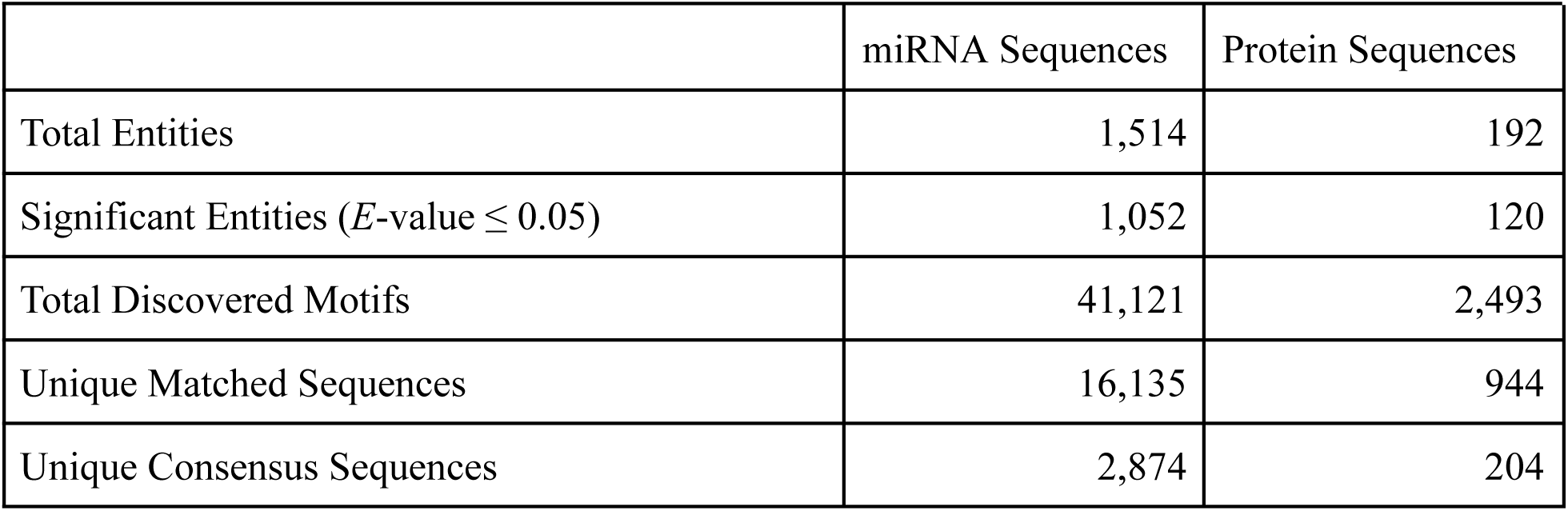
Statistics of discovered motifs from MEME-suite analysis.

The consensus sequences identified in both miRNAs and proteins were paired according to interacting miRNA-protein for molecular docking analysis to assess their binding affinity. The matching sequences against the consensus sequences were extracted from the three-dimensional structure of miRNAs and proteins.

### 3.4. Molecular docking ranked interacting patterns among miRNA-protein interactions

Molecular docking analysis using HDock predicted 100 conformations for the paired motif fragments of the interacting protein-miRNA and ranked using composite index method to yield final ranking of the conformations. Out of the 100 ranked conformations, top ranking conformation was selected for every interacting pair yielding a total of 7,589 interactions among 192 proteins and 824 miRNAs (Supplementary File 7).

Among these interactions, 1,011 interactions were found exclusively in EVs while 273 were exclusive to extracellular environments. This classification was made based on the proteins marked to be associated with EVs and plasma. The most frequently occurring consensus patterns were filtered. We found ALAPPAIP and AAAGUAAUUGY for protein and miRNA, respectively, were the most common consensus sequences (Table 2 and Supplementary File 7).

**Table 2:**
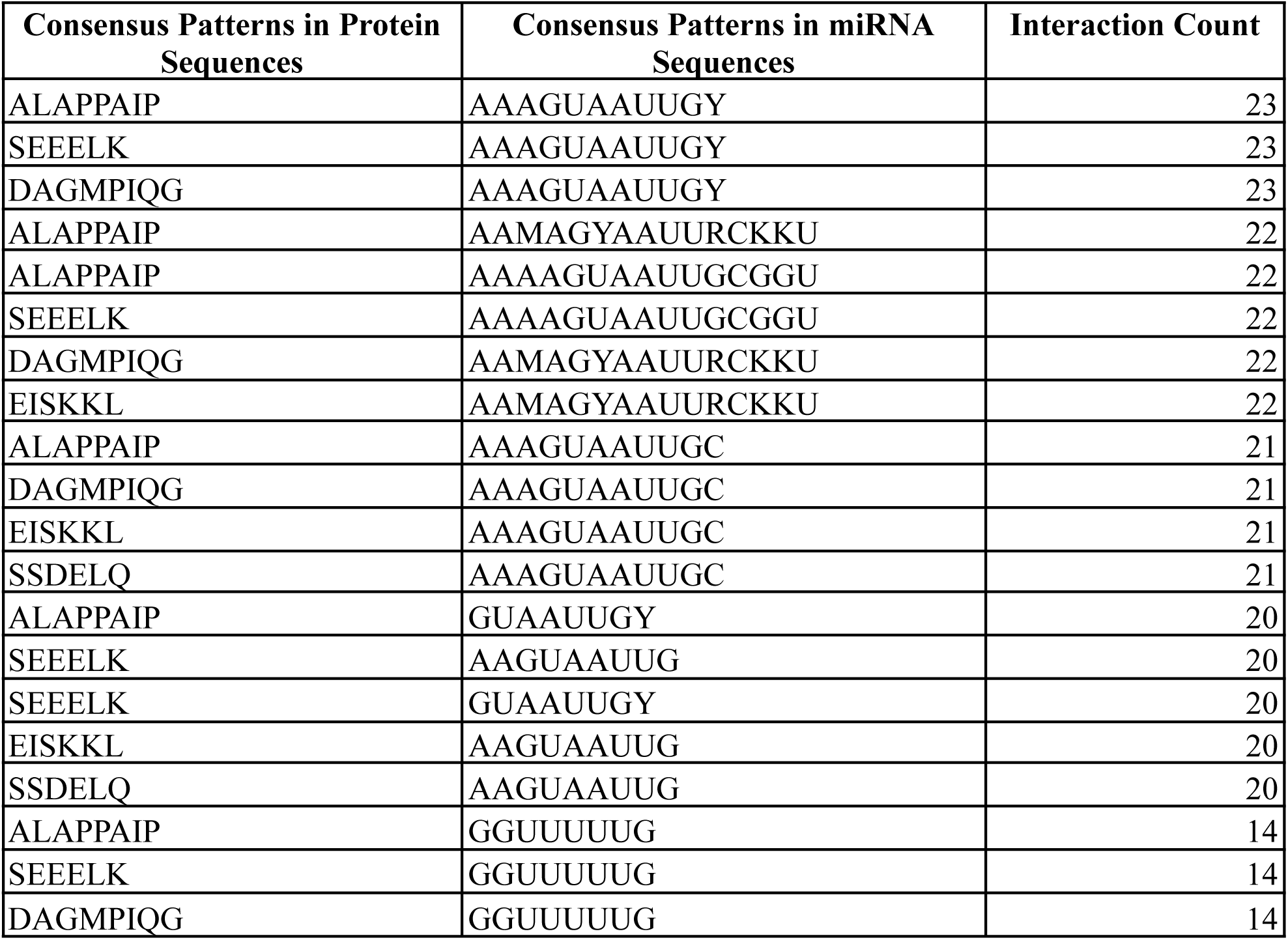
Top 20 interacting patterns among 7,589 miRNA-protein sorted according to their interaction frequency.

### 3.5. Implication of ranked miRNA-protein interaction pattern and correlation with their expression results

Further, the analysis attempted to correlate the interacting miRNA-protein with existing research to complement their findings. The correlations observed between miRNA and protein interacting patterns in cell-free environments suggest that the interactions occurring in the extracellular environment are of utmost importance along with the intracellular counterparts.

#### 3.5.1. Top docking result correlates with findings of neurodegenerative disorders

In our study, the docking of Apolipoprotein B mRNA editing enzyme catalytic subunit 3C (APOBEC3C) and hsa-mir-6511a-5p ranked at top 1 (Table 3 & Supplementary File 7) and it was categorized as the miRNA-protein interaction in extracellular vesicles. Habib et. al. found multiple proteins and miRNAs to be differentially expressed in their study for neurodegenerative disorder [57]. We found an overlap with Habib et al.’s findings that included proteins such as APOBEC3C, Non-POU domain-containing octamer-binding protein (NONO), and Zinc finger CCCH domain-containing protein (ZC3H7B), and miRNAs such as hsa-mir-6511a-5p, hsa-mir-185-3p, hsa-mir-665, hsa-mir-4692, and hsa-mir-6813-5p (Table 3). In another study, Seki et al. found hsa-miR-6813-3p to be a potential serum biomarker in major depression [58]. The interaction and expression correlation can be exploited further to understand the synergistic effect of the network of miRNA-protein interaction to understand the mechanism of progression of neurological disorder and precise characterisation of its stages.

**Table 3:**
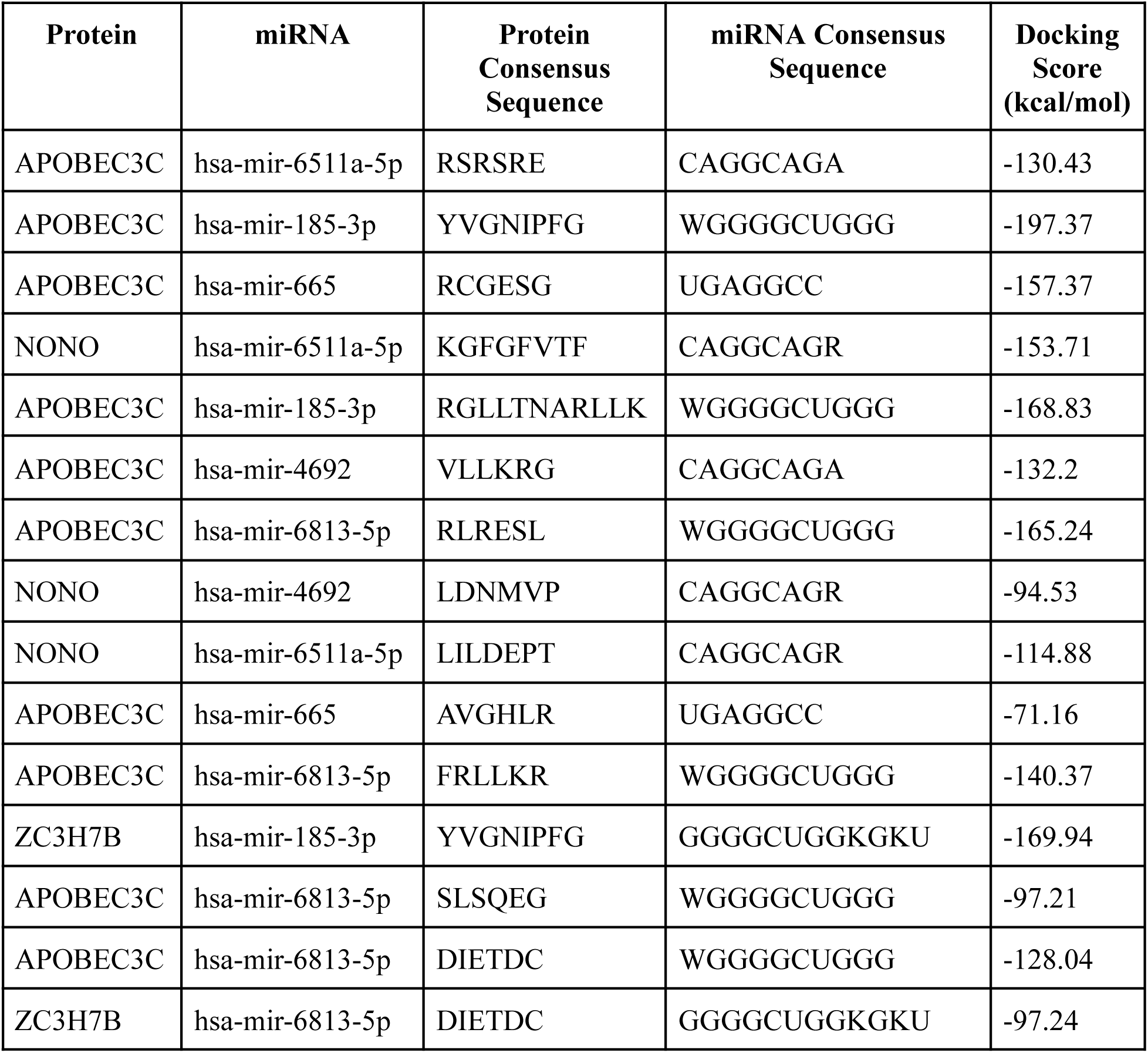
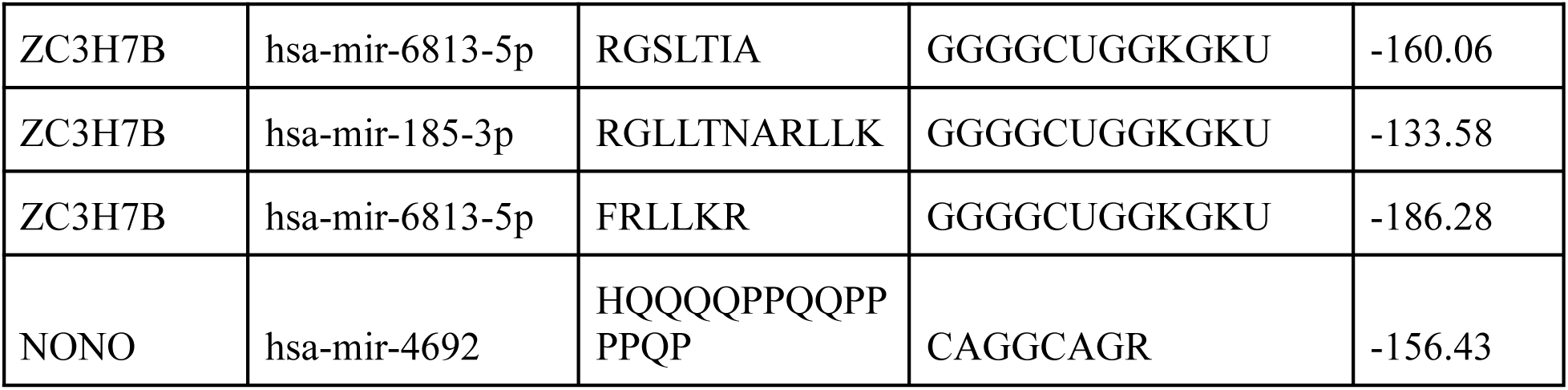
Docking score and interacting consensus sequences of cell-free proteins and miRNAs differentially expressed in neurodegenerative disorders.

The differentially expressed miRNA-protein (Figure 4) having suggested interactions of 15 protein and 5 miRNA consensus sequences (Table 3) can be exploited as therapeutic targets after thorough *in silico* and *in vitro* investigation. The interaction in the extracellular environments can be targeted using small molecules or natural compounds available through diet.

**Figure 4:**
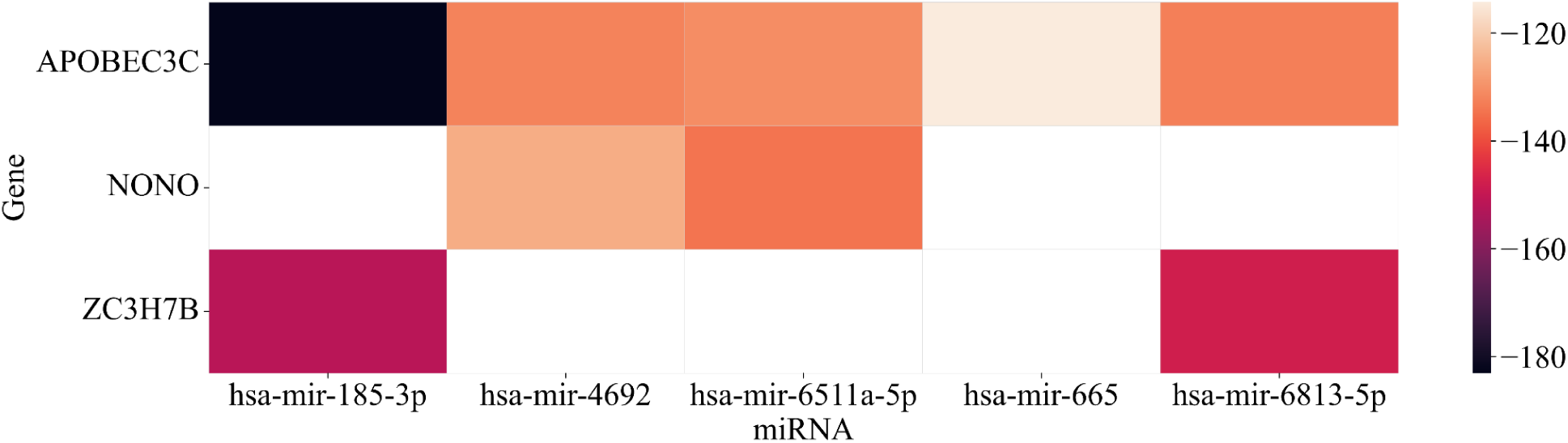
Heatmap plot for the interaction of APOBEC3C, NONO, and ZC3H7B with miRNAs, hsa-mir-185-3p, hsa-mir-4692, hsa-mir-6511a-5p, hsa-mir-665, hsa-mir-6813-5p. Color bar represents the docking score of the interaction where white color shows absence of the interaction.

#### 3.5.2. Potential biomarkers for Schizophrenia extend their interactions with other cf-miRNAs and proteins

Serine and Arginine rich splicing factor 7 (SRSF7) and hsa-miR-4429 are identified as potential biomarkers for Schizophrenia. Davarinejad et al., in another study found that SRSF7 hsa-miR-574-5P, -1827, and -4429 were identified as differentially regulated and among potential biomarkers for Schizophrenia [18]. Our study discovered SRSF7 found to be interacting with hsa-miR-4429. In SRSF7 protein, four sequences were identified to be interacting with the miRNA-4429 *viz.,* YGRYGG, VEFEDP, GLDGKVIC, and SRSRSRSRGRRSRSA with CUGGGAKGDG.

The interaction between SRSF7 and hsa-miR-4429, along with the identified interacting sequences, highlights the intricate regulatory mechanisms involved in gene expression. Further investigation into the functional consequences of this interaction could elucidate its role in cellular processes and disease pathways involved in schizophrenia.

#### 3.5.3. The cf-miRNAs and proteins might carry regulatory machinery for cancer progression and cardiovascular disorders

Our study observed a methyltransferase, N6-adenosine-methyltransferase catalytic subunit (METTL3) in the extracellular group binds to hsa-mir-126-5p, hsa-mir-93-3p, hsa-mir-302a-3p, hsa-mir-33a-3p, and hsa-mir-20b-3p in free from or encapsulated within extracellular vesicles. Additionally it shares its interacting partners with an Argonaute complex (AGO2) (Table 4). Bi et al. reported METTL3 mediating maturation of miR-126-5p leading to ovarian cancer progression via PTEN-mediated PI3K/Akt/mTOR pathway [59]. Xia et al. reported METTL3 mediated extracellular miR-93-5p to be involved in smoking-induced emphysema [60]. Another finding revealed hsa-miR-33a-3p regulating METTL3-mediated inhibition of pancreatic cancer invasion and metastasis by modulation of Epithelial-mesenchymal transitions [61].

**Table 4:**
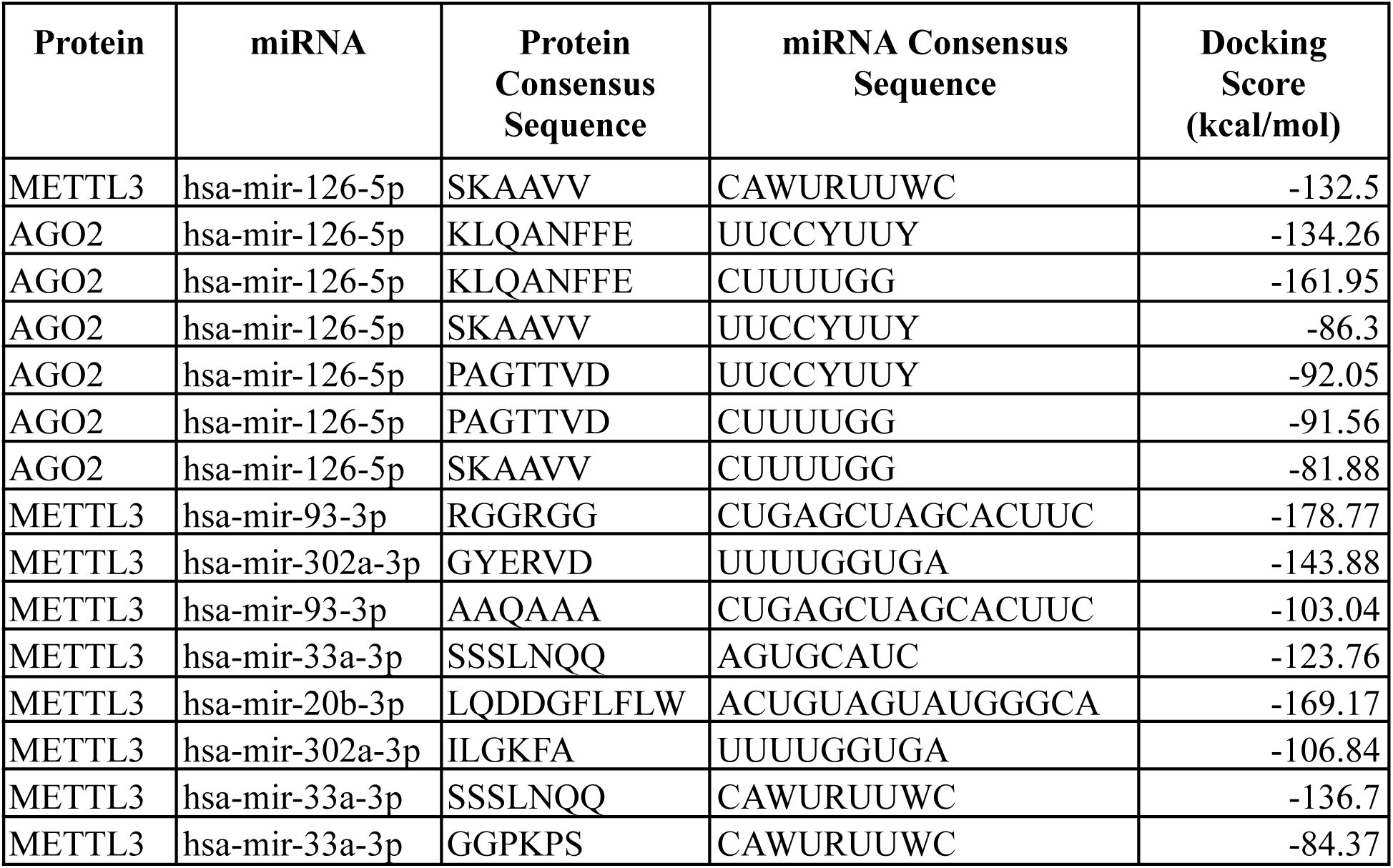
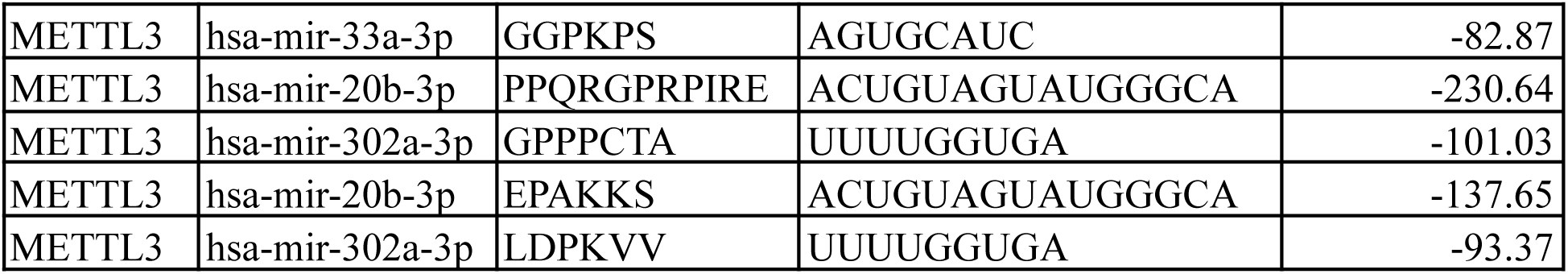
Interaction of METTL3 and AGO2 proteins with miRNAs specially hsa-miR-126-5p: Docking score and consensus patterns.

The complex interplay of AGO2 and METTL3 with different miRNAs may lead to multiple pathways followed by cellular processes to achieve similar phenotypic effect. This intricate interaction network present in extracellular environments can be further explored as a reason for therapy relapse, disease progression biomarkers, and precision therapeutic targets.

Further METTLE3 is also involved in N6-methyladenosine (m6A) modification of eukaryotic RNAs along with many other proteins and this modification is detected by m6A-binding proteins. We found IGF2BP1, HNRNPA2B1, and YTHDF1 in our study that could detect m6A modification and also show their interaction in an extracellular environment. The m6A methylation is widely studied in relation with human ischemic stroke. Further, a natural plant based potent antioxidant flavonol, Quercetin, is reported to inhibit METTL3 and also regulates miR-26b, mIR-320a, and miR-126 miRNAs in different studies. Quercetin targets m6A modification to inhibit cancer cell proliferation [62].

We speculate that the interactions between METTL3 and miRNAs (Table 4) could be targeted by different natural compounds that might show different affinity for different METTL3-miRNA interactions at different stages of diseases like different types of cancers or cardiovascular disorders.

#### 3.5.4. Protein modification site uphill a target motif

While attempting to discover existing relationships between motifs and interacting entities along with their significance of newly discovered motifs through literature review and various databases, the searches yielded no satisfactory results except a few. In a study, Hanoulle et al. identified a site that had sequence SEEELK downstream the modification site in the study to identify purine nucleotide binding sites in complex proteomes [63]. In our study, the motif was discovered in PTBP1, XRCC6, SDAD1, and G3BP1 and was among top motifs (Table 2) however the study reports protein modification sites only in XRCC6. The pattern interacted with 22 miRNAs belonging to the hsa-miR-548 family with seven unique miRNA consensus sequences and one miRNA hsa-miR-559 with sequence AAAGUAAUUGY (Figure 5).

**Figure 5:**
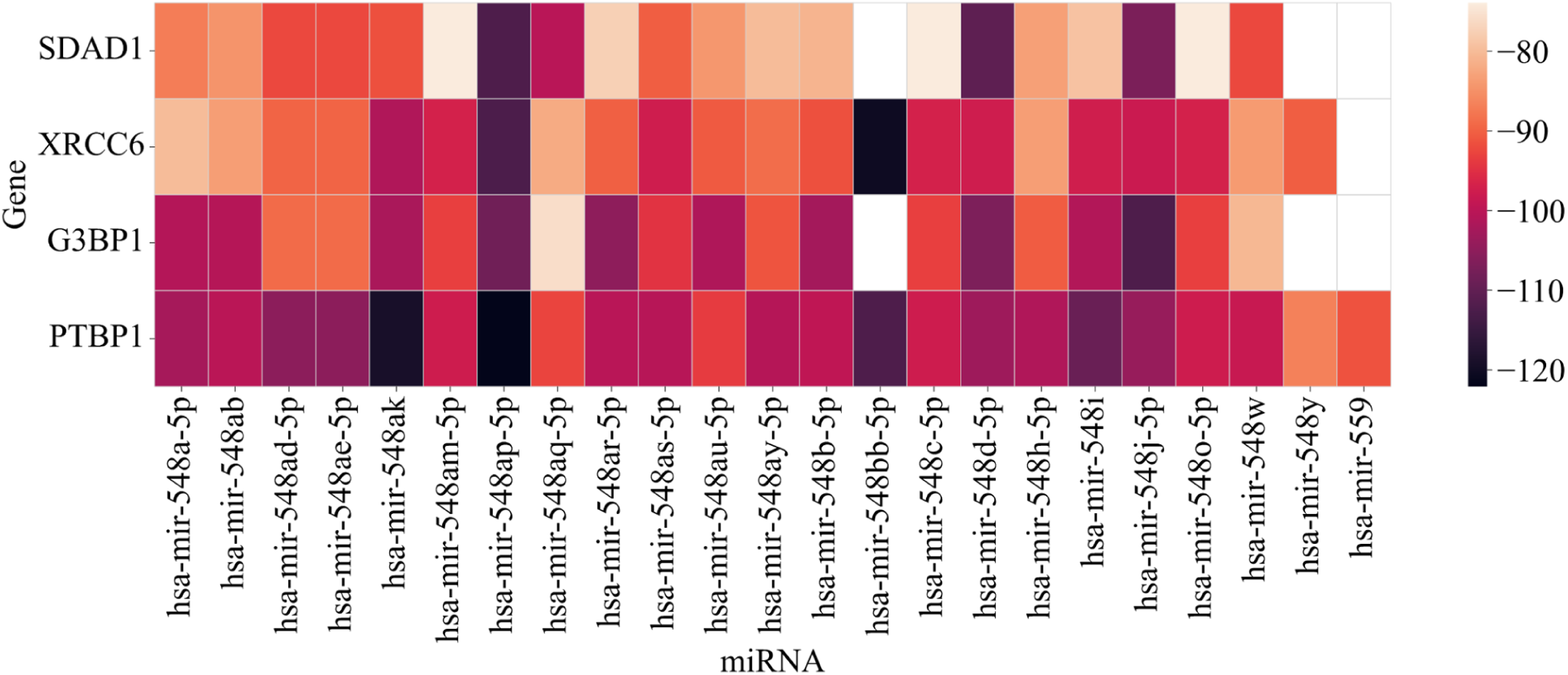
Heatmap plot miRNA-protein interactions where miRNAs interact with consensus sequence SEEELK. Color bar represents interaction strength in terms of docking score The information of similar interactions of multiple miRNAs with a possible modification site could be further exploited to understand the mechanism of miRNA binding with protein to regulate cellular homeostasis.

#### 3.5.5. MiRNAs belonging to hsa-mir-548 family could be baseline miRNAs participating in multiple diseases

Different members of the family, hsa-miR-548, are differentially expressed in different disease types and have been attributed for their association with disease phenotypes including metastasis promotion, therapeutic target in human breast cancer, diabetic retinopathy biomarker, and differential regulation in age-related neurodegenerative diseases [2,24,64,65]. Further, exploring expression of miR-548 family members using CancerMIRNome it was found that hsa-miR-548a-3p is overexpressed in different cancer types like benign ovarian disease, ovarian cancer, breast cancer, pancreatic cancer, esophageal cancer, colorectal cancer, hepatocellular carcinoma, gastric cancer, and lung cancer [66].

This hsa-mir-548 microRNA family members (total 70 members overlapping our study) yielded the highest interactions as they interacted with 43 proteins (Figure 6). These interactions involved 131 protein consensus sequences while only 51 consensus sequences from miRNAs. This interactome has relatively similar interactions yet is involved in the regulation of diverse biological processes.

**Figure 6:**
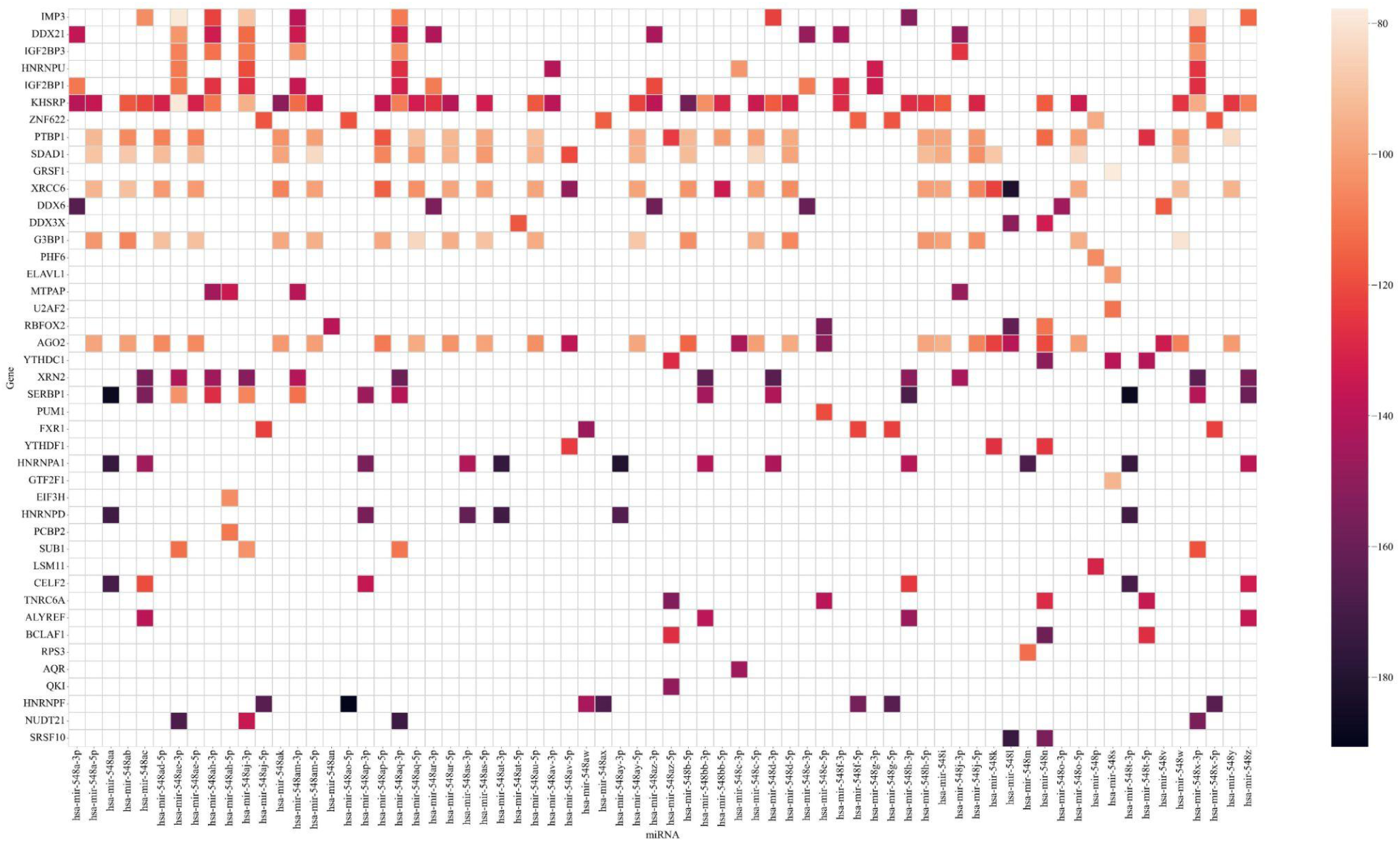
Heatmap plot for miR-548 family members with proteins in extracellular environment. The color bar represents docking score of the interactions The miRNA-548 family miRNAs and their target proteins participate in varied biological processes yet share predictable similar interactions, due to this characteristic miRNA-548 family members might act as a model to understand the sequence based interactions of miRNA-protein or other types of interactions like miRNA, mRNA, DNA, or protein occurring in extracellular environment or within the cell.

#### 3.5.6. miR-30 family regulates mitochondrial fission

hsa-miR-30 family miRNAs could interact with the proteins CPEB4, CPSF1, CPSF3, CPSF4, FXR1, LARP4, LIN28B, MBNL2, RBM5, STAU1, TNRC6A, YTHDC1, and YWHAG genes along with other reported target genes such as p53. Being a valuable biomarker, hsa-miR-30 family members exert their role in mitochondrial fission by directly interacting with cellular p53 [67,68]. Almost 11 unique members interact with 30 proteins with their 56 consensus sequences. Interestingly, the hsa-miR-30 family is involved in the development and diseases of bone tissue, reproductive system, adipogenesis, and other normal tissues and organs [19,21,67,69,70].

There are numerous miRNA-protein combinatorial interactions that have significant importance and have been partially correlated. Exploring the importance of each miRNA-protein interaction is out of the scope of this study therefore it limits findings to few results from literature review. These findings can be explored further using *in silico*-based for drug discovery and machine learning algorithms to provide insight into the lifecycle of miRNA and its interacting partners. The outcomes will be easy to translate into theranostic application as the possible pathway of interacting entities will be known along with the consequences of targeting them.

## 4. Conclusion

The correlation of miRNA pool with protein in the context of normal condition and disease condition is gaining importance to have a better purview of precise mechanisms involved in the disease onset and progression. Our study finds new sequence patterns that are not widely reported along with miRNA protein interactions that have not been discovered yet can be significant in precise disease diagnosis and therapeutic target discovery.

We strongly recommend therapeutic research involving primary, secondary and tertiary structure based interaction between different biomolecules whether it is intracellular or extracellular. In our study we have discovered possible motifs of miRNA and proteins that can be transitioned to aid the experimental findings in a disease context. This further strengthens the findings that direct miRNA-protein interactions in extracellular space should be investigated along with the intracellular interactions for precise diagnosis.

The pool of carrier proteins in the blood plasma sharing consensus sequences to interact with miRNAs can be employed to look into alternate pathways when there is therapy relapse, drug resistance, or the drugs show different effects in different individuals. We suggest that the exploration of the pool of AGO2 and METTL3 interaction and miR-548 and miR-30 families in extracellular space should be prioritized in the context of disease as the miRNA family holds similar structure and interaction pattern but have varying phenotypic effects. A disease is not regulated by a single protein or miRNA but is a synchronized effect of interactions of different biomolecular species and miRNA or protein are prominent biomolecules controlling the gene regulation and phenotypic effect. Therefore, therapeutic approaches require to consider a wide range of interactions to overcome the disease relapse, drug inefficacy, or drug resistance.

Availability of a larger interactome and evidence of interactions of miRNA-protein has limited our study to find the possible patterns and implication of these interactions. This gives significant weightage that the specific interactions can be categorized in a common way compared to normal conditions and in disease-specific cases. Additionally suggesting drugs targeting these sequence based interactions was out of scope of our study. As our understanding of the structural complexities of miRNA-protein interactions deepens, innovative therapeutic approaches, such as small molecules or RNA-based therapeutics, can be designed to modulate these interactions for precise and effective disease management in extracellular environment rather than approaching to deliver drugs intracellularly.

In summary, miRNA-protein motif interactions are crucial for regulating gene expression and influencing various biological processes. Targeting these interactions holds promise for developing novel therapeutic strategies for cardiovascular disease, neurological disorders, disorders of the immune system, and other chronic diseases like cancer. Thus, unraveling the complexities of miRNA-protein motif interactions not only enhances our comprehension of cellular regulatory networks but also opens new avenues for the development of targeted therapeutics with broader implications for human health.

## Supporting information

SF1-Proteins

SF2-miRNAs

SF3-Extracellular-miRNA-Protein-Interactions

SF4-DAVID-GSEA-Protein

SF5-DAVID-GSEA-miRNA

SF6-Discovered-Motifs

SF7-Molecular-Docking-Results

## Acknowledgements

Authors acknowledge Dr. D. Y. Patil Biotechnology and Bioinformatics Institute, Pune, India for research infrastructure and the Department of Science & Technology (DST), Government of India for providing DST-FIST grant (SR/FST/LS-I/2017/70) to support research infrastructure and instruments. VKS acknowledges the Council of Scientific and Industrial Research (CSIR), New Delhi, India for CSIR-SRF fellowship (File No.: 09/1340(11487)/2021-EMR-I). SS acknowledges Department of Biotechnology, Government of India for Ramalingaswami Re-entry Fellowship grant (File No.: BT/RLF/Re-entry/47/2021). SB acknowledges intramural grant Intramural of Dr. D. Y. Patil Vidyapeeth (DPU), Pune, India (DPU/644-43/2021).

## Supplementary Materials

Supplementary File 1: Table of extracellular proteins

Supplementary File 2: Table of extracellular miRNAs

Supplementary File 3: Table of extracellular interactions

Supplementary File 4: Table of Protein GSEA

Supplementary File 5: Table of miRNA GSEA

Supplementary File 6: Table of discovered MEME motifs

Supplementary File 7: Molecular docking results

